# A bioprinted silk marrow niche reveals mechanical regulation of human megakaryopoiesis under genotoxic stress

**DOI:** 10.64898/2026.06.18.733176

**Authors:** Christian A. Di Buduo, Matteo Migliavacca, Michele Menotta, Elin Pernevik, Alessandro Malara, Santo Diprima, Giovanni Goggi, Claudia Del Fante, Monica Savio, Volodymyr Kuzmenko, Mauro Magnani, Alessandra Balduini

**Affiliations:** Department of Molecular Medicine, University of Pavia, Pavia, Italy; Silk4B s.r.l., Milan, Italy; Department of Biomolecular Sciences, University of Urbino “Carlo Bo”, Urbino, Italy; CELLINK Bioprinting AB, Gothenburg, Sweden; Center for Omics Sciences, IRCCS San Raffaele Scientific Institute, Milan, Italy; Division of Immunohaematology and Transfusion Service, Fondazione IRCCS Policlinico San Matteo, Pavia, Italy; Department of Biology and Biotechnology, University of Pavia, Pavia, Italy

## Abstract

Hematopoietic stem and progenitor cells (HSPCs) reside in a mechanically distinct bone marrow niche, yet how niche biomechanics shape genome stability and stress responses has been difficult to test because conventional two-dimensional (2D) culture lacks marrow viscoelasticity and uses surfaces that activate platelets, confounding hematopoietic readouts. Here, we show that this methodological gap has masked a basic principle: the marrow niche actively constrains genotoxic stress signaling in HSPCs, and 2D culture systematically overstates DNA damage and impairs differentiation *in vitro*. We engineered silk fibroin, a biologically inert biomaterial that does not activate platelets and recapitulates marrow viscoelasticity, into SilkInk, a 3D-bioprintable bioink, and used it to reconstruct a biomimetic marrow microenvironment. HSPCs encapsulated in SilkInk preserved clonogenic potential and multilineage differentiation, whereas 2D-cultured HSPCs activated cytoskeletal-tension and genome-surveillance programs characteristic of chronic stress, including pathways related to replication stress, DNA damage response, and redox stress. Cell phenotyping and single-cell RNA sequencing during megakaryopoiesis revealed that SilkInk supported coordinated endomitotic progression and terminal maturation, with progression from CD34^+^CD61^−^CD41^−^CD42b^−^ progenitors to CD34^−^CD61^+^CD41^+^CD42b^+^ megakaryocytes, including increased 8N and >16N populations, whereas 2D culture and conventional 3D hydrogels sustained DNA damage signaling and impaired thrombopoiesis. The same hierarchy held under cytotoxic challenge, as 5-fluorouracil amplified DNA damage and crippled platelet output in 2D, whereas SilkInk-encapsulated HSPCs maintained differentiation, mirroring native marrow resilience. These findings reposition niche mechanics as an active determinant of hematopoietic genome stability and establish SilkInk as a physiologically faithful platform for studying hematopoiesis and predicting marrow responses to chemotherapy.

## Introduction

The bone marrow microenvironment regulates hematopoiesis through an integrated set of biochemical, structural, and mechanical cues that coordinate HSPC maintenance, lineage commitment, and stress adaptation ^1–3^. Dysregulation of these inputs underlies hematologic disorders, bone marrow failure, and treatment-induced cytopenias, particularly when genome instability or replicative stress is involved. Megakaryopoiesis is especially sensitive: lineage maturation depends on endomitosis and tightly controlled DNA replication dynamics, and defects in DNA damage response (DDR) pathways are increasingly recognized as central drivers of impaired thrombopoiesis in thrombocytopenia, myeloproliferative neoplasms, and chemotherapy-induced marrow toxicity ^4–9^. How hematopoietic cells engage these stress programs, and how the niche shapes that engagement, therefore demands experimental systems that faithfully reproduce the native bone marrow microenvironment.

Most *in vitro* hematopoiesis studies still use conventional two-dimensional (2D) culture, which fails to recapitulate the bone marrow’s three-dimensional (3D) architecture, viscoelasticity, and spatial organization ^10,11^. Removed from their niche, HSPCs rapidly lose stem-cell function and activate stress-associated programs that distort cell identity and differentiation potential ^12,13^. Conventional substrates actively induce oxidative stress, replication-associated DNA damage, and proteostatic imbalance, raising the possibility that much of what is interpreted as “intrinsic” HSPC stress *in vitro* is partly an artifact of substrate mechanics ^13–26^. These distortions are especially consequential for megakaryocyte differentiation, where standard systems poorly support the maturation trajectories required for proplatelet formation and platelet release ^27–29^.

Bioengineered bone marrow models, organ-on-chip systems, 3D scaffolds, and iPSC-derived organoids have made important progress in reconstructing aspects of marrow organization and multicellular interaction ^27,30–40^. Yet most remain limited by low reproducibility, complex culture requirements, modest scalability, or dependence on heterogeneous extracellular matrices and supporting stromal populations. A simplified, mechanically defined system capable of sustaining physiologically faithful hematopoietic programs, including megakaryocyte maturation and thrombopoiesis, has been lacking.

Silk fibroin from *Bombyx mori* meets these requirements^41–43^. It forms stable networks with tunable mechanics and intrinsic antioxidant activity, and it preserves and presents bioactive molecules under mild aqueous processing ^43–48^. Critically for hematopoietic applications, silk fibroin is biologically inert and does not activate platelets ^41^, a property that few alternative biomaterials offer and that is indispensable for modeling megakaryopoiesis and thrombopoiesis *in vitro*.

Here, we developed SilkInk, a silk-based bioink optimized for reproducible 3D bioprinting of biomimetic marrow niches, and combined proteomic profiling, single-cell RNA sequencing, and functional assays to ask how microenvironmental context regulates HSPC maintenance, megakaryocyte maturation, and stress responses. We find that conventional 2D culture and standard 3D hydrogel bioinks induce sustained activation of cytoskeletal-tension and genome-surveillance programs that impair differentiation and thrombopoiesis, whereas SilkInk suppresses these programs and supports orderly megakaryocyte maturation. Under 5-fluorouracil challenge, SilkInk-encapsulated HSPCs preserve differentiation and recapitulate adaptive responses consistent with native marrow resilience, while 2D culture amplifies DNA damage and collapses platelet output. Within this system, thrombopoietin receptor agonists (TPO-RAs) rescue stress-impaired thrombopoiesis, resolving injury, adaptation, and recovery into a unified trajectory.

## Materials and Methods

### Preparation of SilkInk and bioprinting parameters

Silk fibroin aqueous solution was obtained from *Bombyx mori*, as previously described ^49^. SilkInk was prepared and bioprinted, as previously described ^47^, using a BIO-X extrusion bioprinter (Cellink Bioprinting). A thermo-regulated printhead enabled precise temperature control over the needle, ensuring a uniform temperature down to the print surface. After bioprinting, the scaffold was crosslinked for at least 15 minutes in a physiological salt solution containing CaCl_2_. Then, the 3D construct was immersed in cell culture medium and kept at 37 °C and 5% CO_2_.

### Cell bioprinting and culture

Human blood samples were obtained from healthy controls after informed consent. All samples were processed in accordance with the ethical committee of the I.R.C.C.S. Policlinico San Matteo Foundation and the principles of the Declaration of Helsinki. CD34^+^ cells were separated by an immunomagnetic bead selection kit (Miltenyi Biotec), as previously described ^50,51^. Samples were cultured at 37 °C in a 5% CO_2_-humidified atmosphere in StemSpan medium (StemCell Technologies). For HSPC maintenance, samples were cultured with 1% penicillin-streptomycin (P/S), 1% L-glutamine, Stem Cell Factor (SCF) 100 ng/mL, FMS-like tyrosine kinase 3 ligand (FLT3-L) 100 ng/mL, Thrombopoietin (TPO) 20 ng/mL, and Interleukin (IL)-6 20 ng/mL. For megakaryocyte differentiation, samples were supplemented with 1% P/S, 1% L-glutamine, 10 ng/mL TPO, and 10 ng/mL IL-11. All cytokines were from Peprotech.

HSPCs or megakaryocyte progenitors were mixed with SilkInk and heated to 37 °C before printing, as described above. In some experiments, cells were mixed into CELLINK RGD bioink (Cellink Bioprinting) and 3D bioprinted according to the manufacturer’s instructions. For SilkInk dissolution, samples were incubated in a formulation comprising sodium citrate, dissolved in a physiologic salt solution. Samples were incubated at 37 °C to dissolve/recover cells from SilkInk. For the Colony-Forming Unit (CFU) assay, a total of 2×10^3^ cells were mixed with MethoCult™ H4434 Classic (STEMCELL Technologies) according to the manufacturer’s instructions. CFUs were manually quantified and classified on day 14 post-seeding under an optical microscope.

### Statistics

Values were expressed as mean plus or minus the standard deviation (mean ± SD), or median and range. The Student’s t-test or ANOVA, followed by a Bonferroni posttest, was used to analyze the experiments. A value of at least p < 0.05 was considered statistically significant.

Additional materials and methods are described in the ‘*Supplemental File*’. Data supporting the results of this study are available within the study. All data sets generated are available on request from the corresponding author, Prof. Alessandra Balduini.

## Results

### SilkInk preserves a primitive HSPC phenotype that 2D culture rapidly erodes

We engineered an extrusion-compatible silk-based bioink, hereafter referred to as SilkInk, for standardized 3D bioprinting of human bone marrow niches and compared its ability to support hematopoiesis with that of 2D standard culture plates (**Figure 1A**). The SilkInk formulation combines regenerated *Bombyx mori* silk fibroin with gelatin, which provides a thermo-reversible sol-gel transition for printability, and sodium alginate, which enables rapid ionic crosslinking ^47^. Extrusion through a 20G nozzle at 8-10 mm s^-1^ provided an optimal compromise between printing fidelity and accurate material deposition, yielding multilayered constructs of a ‘*flower*’-shaped geometry (**Figure 1A**). Human CD34^+^ HSPCs embedded in SilkInk were distributed evenly throughout the construct volume and across depth, confirming that bioprinting generates spatially uniform 3D hematopoietic microenvironments (**Figure 1B**).

**FIGURE 1.**
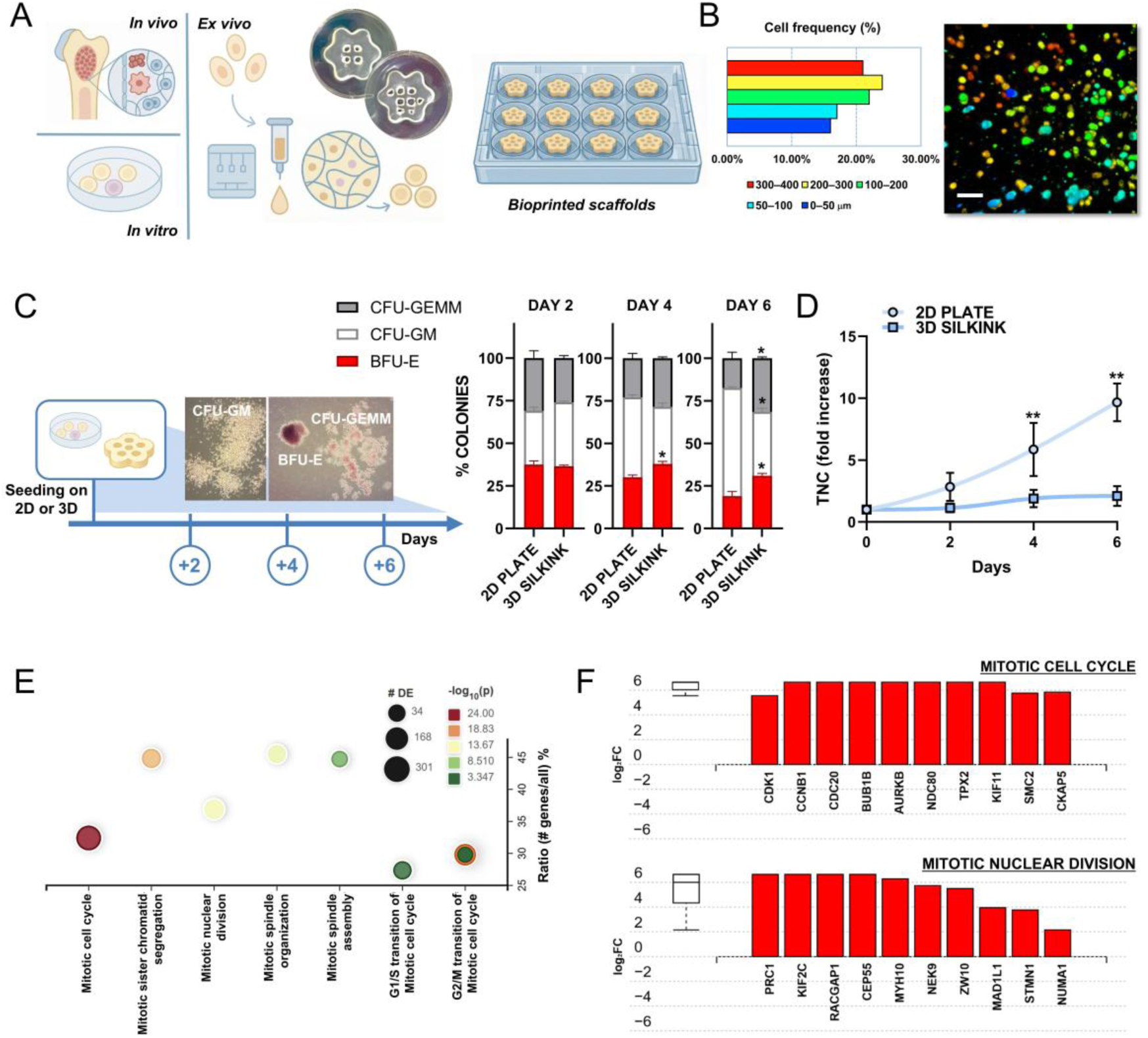
3D silk niches support HSPC culture and preserve clonogenic potential. (A) Schematic overview of the experimental workflow, from the native bone marrow niche to *in vitro* culture, and *ex vivo* generation of bioprinted 3D scaffolds. (B) Left, depth distribution of cells across the printed construct, reported as the percentage of cells detected within the indicated z-intervals (0-50, 50-100, 100-200, 200-300, and 300-400 µm). Right, representative fluorescence image used for cell localization analysis. Cells were stained for CD34 and presented as pseudocolors indicating their position along the z-axis. Scale bar, 50 µm. (C) Experimental layout for colony-forming unit (CFU) analysis after seeding primary human HSPCs in 2D plates *versus* 3D SilkInk and retrieving cells after 2, 4, or 6 days; representative images of CFU-GM, CFU-GEMM, and BFU-E colonies are shown. The stacked bars report the relative distribution of colony types at each time point. *p < 0.05 (D) Total nucleated cell (TNC) expansion over time in 2D plates and 3D SilkInk culture, expressed as fold increase relative to day 0. **p < 0.01 (E) Functional enrichment dot plot highlighting mitotic and cell-cycle-related biological processes differentially represented between 2D and 3D conditions. Dot size indicates the number of proteins, and dot color indicates significance. (F) List of proteins contributing to the enrichment of the indicated mitotic cell-cycle and mitotic nuclear division programs. Data are shown as mean ± SD.

The 3D niche preserved a broader hematopoietic output than conventional 2D culture plates. CFU assay performed using HSPCs retrieved after 2, 4, and 6 days of culture showed that cells from SilkInk sustained CFU-GEMM, CFU-GM, and BFU-E output across the time course, whereas 2D cultures progressively collapsed toward CFU-GM dominance (**Figure 1C**). Total nucleated cell counts increased markedly in 2D compared with SilkInk (**Figure 1D**), indicating that 2D expansion comes at the cost of multilineage potential. Proteomic analysis explained this trade-off: in 2D, the most enriched processes were mitotic cell cycle and nuclear division programs (**Figure 1E**), with elevated levels of core mitotic regulators (CDK1, CCNB1, CDC20, BUB1B, AURKB, NDC80, TPX2, KIF11, SMC2, and CKAP5), and mitotic nuclear division factors (PRC1, KIF2C, RACGAP1, CEP55, MYH10, NEK9, ZW10, MAD1L1, STMN1, and NUMA1, among others; **Figure 1F**). 2D culture, therefore, drives HSPCs into a proliferation-biased state at the expense of stemness, whereas SilkInk preserves a phenotype closer to the more primitive HSPC compartment.

### 2D culture imposes replication stress and genome surveillance programs on human HSPCs

Volcano-plot analysis of 2D *versus* 3D proteomes revealed that HSPCs from 2D cultures preferentially accumulated proteins involved in cytoskeletal tension, nuclear force transmission, and stress signaling, including ROCK1, ROCK2, SRC, SUN1, and SYNE2 (**Figure 2A**). Other cytoskeletal regulators (MYO1D, ACTG2, ARHGAP26) were concomitantly reduced, indicating that 2D imposes selective remodeling of Rho-dependent mechanical networks rather than generalized cytoskeletal upregulation. The mechanical environment of SilkInk supports this interpretation. Across production batches, SilkInk showed shear-thinning behavior, viscosity recovery after repeated high-shear cycles, and a temperature-dependent viscoelastic transition (**Supplemental Figures 1A-C**). After printing and culture at 37 °C, scaffolds exhibited reproducible viscoelastic and compressive properties (**Figures 2B and 2C; Supplemental Figures 1D and 1E**), with stiffness in the low-kPa range, close to native marrow compliance, while tissue-culture plastic is orders of magnitude stiffer (**Figure 2D**). SilkInk, therefore, provides HSPCs with mechanical inputs that approximate their *in vivo* microenvironment ^3^, whereas 2D culture does not.

**FIGURE 2.**
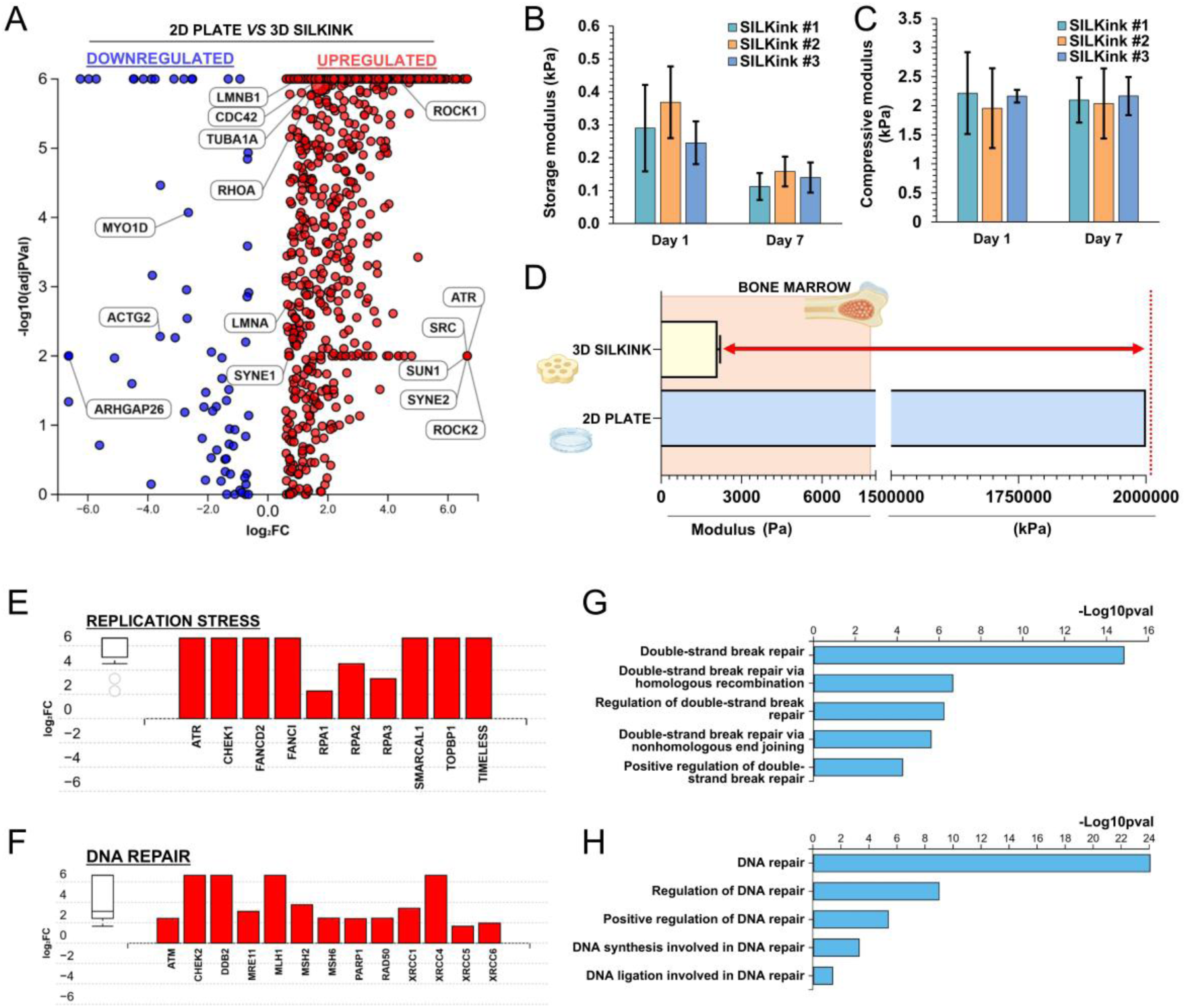
Proteomic features of HSPCs in SilkInk *versus* 2D culture plates. (A) Volcano plot showing differentially expressed proteins in 2D plates *versus* 3D SilkInk cultures. Selected proteins are annotated. (B) Storage modulus of three independent SilkInk batches measured at day 1 and day 7 post-printing. (C) Compressive modulus of the same independent SilkInk batches. (D) Comparison of the stiffness of the 3D SilkInk construct and conventional 2D culture plates, with native bone marrow reference range. Representative proteins associated with replication stress (E) and DNA repair (F) programs, enriched in 2D *versus* 3D culture. (G, H) Ontology enrichment analyses of DNA repair, DSB repair, and genome-maintenance programs. Data are shown as mean ± SD.

These mechanical differences mapped onto distinct molecular states. Biological Process and Hallmark enrichment analyses (**Supplemental Figures 2A and 2B**) showed that proteins upregulated in 2D were enriched for stress-response pathways, with Hallmark sets identifying G2/M checkpoint, DNA repair, and mitotic spindle activation. 2D-cultured HSPCs accumulated canonical replication-stress effectors (ATR, CHEK1, RPA1-3, FANCD2, FANCI, SMARCAL1, TOPBP1, TIMELESS; **Figure 2E**) and DNA repair factors spanning Double-Strand Break (DSB) sensing, Homologous Recombination (HR), and Non-Homologous End Joining (NHEJ) (e.g., ATM, CHEK2, MRE11, RAD50, PARP1, MSH2/6, XRCC1/4/5/6, DDB2; **Figure 2F**). Pathway enrichment analysis confirmed upregulation of DNA repair programs (**Figures 2G and 2H**). Parallel elevation of NADPH oxidase subunits (e.g., NOX1, CYBB, NOX4, CYBA, NCF1/2/4, RAC1, RAC2) and antioxidant enzymes (e.g., HMOX1, NQO1, GCLC, GCLM, SLC7A11, TXNRD1) further indicated a redox-stressed state engaged with cytoprotective responses. Together, these data establish that 2D culture imposes a chronic mechanotransduction and genome-surveillance-driven stress program on HSPCs.

### SilkInk supports coordinated megakaryocyte maturation and higher platelet output

We next investigated how distinct culture conditions influence HSPC differentiation toward megakaryopoiesis. Single-cell transcriptomic analysis identified a clear developmental continuum spanning early progenitors, late progenitors, and mature megakaryocytes, with a small population of exhausted megakaryocytes at the terminal end of the trajectory (**Figures 3A and 3B; Supplemental Figure 3**). Early progenitors expressed lineage-commitment factors (*GATA1, FLI1, NFE2*); late progenitors peaked for proliferation and endomitosis genes (*MKI67, TOP2A, CCNB1, CDC20, PLK1*); and terminal megakaryocytes engaged platelet-biogenesis and cytoplasmic-remodeling programs, including genes for the integrin ITGA2B/ITGB3, the vWF receptor components GP1BB and GP9, the platelet structural gene TUBB1, the α-granule factors PF4 and PPBP, and actin-integrin coupling machinery (**Figure 3C and Supplemental Figure 3**). Cell-cycle inhibitors such as *CDKN2D* emerged in mature compartments, consistent with coordinated cell-cycle exit preceding cytoplasmic maturation and proplatelet formation (**Supplemental Figure 3**). All programs were broadly suppressed in the exhausted megakaryocyte population (**Figure 3C**). A defining feature of this trajectory was a discrete endomitotic phase preceding terminal maturation. Heatmap analysis confirmed that late progenitors were the most enriched compartment for DNA-replication and endomitosis genes, including *CDK1, CDC45, AURKB, CCNB2, NUSAP1, CDCA8, RACGAP1, GINS2, ORC1, MCM4,* and *CHEK1* (**Figure 3D and Supplemental Figure 3**). EdU incorporation tracked the same logic: low in CD61^-^CD41^-^CD42b^-^ early progenitors, peaked in CD61^+^CD41^+^CD42b^-^ late progenitors, and declined in mature CD61^+^CD41^+^CD42b^+^ megakaryocytes (**Figure 3E**). DNA synthesis is, therefore, concentrated within an intermediate stage and downregulated to permit terminal differentiation. Phenotypic analysis confirmed orderly progression from low-ploidy, CD34^+^CD61^-^CD41^-^CD42b^-^progenitor-enriched compartment (early progenitors), to an intermediate CD34^low^CD61^+^CD41^+^CD42b^-^ population (late progenitors), and ultimately polyploid CD34^-^CD61^+^CD41^+^CD42b^+^ megakaryocytes (**Supplemental Figure 4**). During megakaryocytic differentiation, cells maintained in rigid 2D culture upregulated mechanotransduction-related pathways, including the CDC42 and RHO GTPase cycles, particularly in late progenitors and megakaryocytes (**Figure 3F).** This indicates that the 2D environment imposes increased cytoskeletal tension during the stages that precede and accompany thrombopoiesis. Consistent with this interpretation, immunofluorescence analysis revealed that megakaryocytes generated in 2D culture displayed a more rounded morphology, with limited cytoplasmic remodeling and fewer proplatelet-like extensions. In contrast, megakaryocytes differentiated within 3D SilkInk showed more megakaryocytes exhibiting cytoplasmic protrusions and evident platelet-like particle shedding (**Figure 3G**). Quantification confirmed that SilkInk significantly increased the percentage of proplatelet-forming megakaryocytes compared with 2D culture (**Figure 3H**). Treatment of 2D cultures with Y-27632, a selective ROCK inhibitor acting downstream of RHO signaling, partially rescued proplatelet formation, supporting the idea that excessive RHO/ROCK-dependent actomyosin tension limits thrombopoiesis in rigid 2D culture conditions (**Figure 3H**). In line with this, platelet-like particle production was markedly higher in SilkInk than in 2D and was also partially restored in 2D upon Y-27632 treatment (**Figure 3I**). Thus, SilkInk, through its ultra-soft 3D microenvironment, provides optimal conditions to improve cellular behavior and preserve the functional cell phenotype without relying on chemical or pharmacological treatments.

**FIGURE 3.**
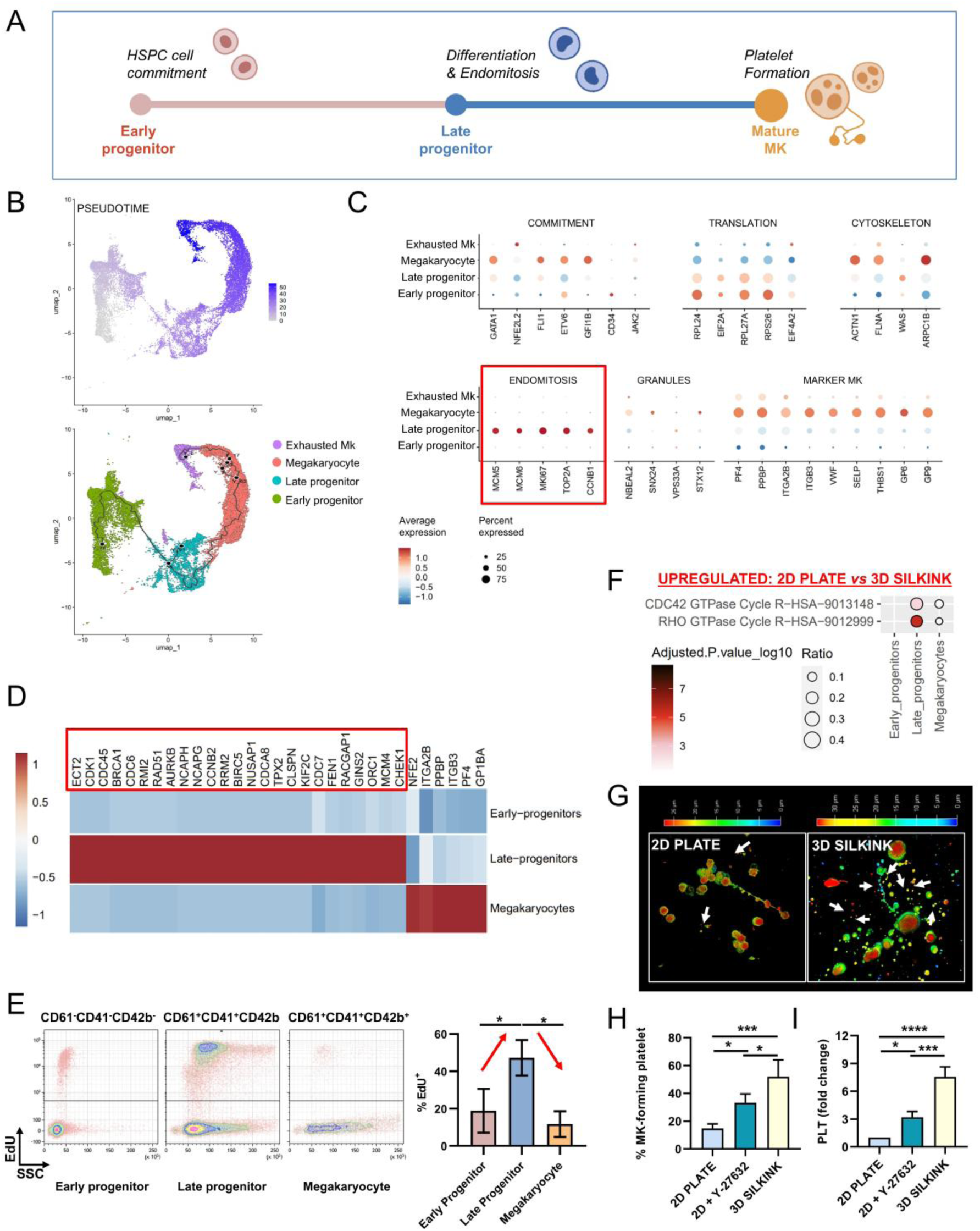
The 3D SilkInk supports improved thrombopoiesis. (A) Schematic representation of megakaryopoiesis, from progression of early progenitors to late progenitors undergoing differentiation and endomitosis, to mature megakaryocytes (MKs), forming platelets. (B) UMAP/pseudotime projection identifying early progenitor, late progenitor, and megakaryocyte states along the differentiation trajectory. (C) Dot plot of representative genes associated with commitment, cytoskeletal remodeling, translation, endomitosis, and mature megakaryocyte identity across the indicated populations. (D) Heat map showing stage-associated gene expression changes from early progenitors to mature megakaryocytes. (E) Representative flow-cytometry plots and quantification of EdU incorporation in early progenitors, late progenitors, and megakaryocytes, showing the highest proliferative/DNA synthesis activity in late progenitors. (F) Dot plot showing the ontology/pathway terms upregulated in the 2D plate compared with the 3D SilkInk culture across early progenitors, late progenitors, and megakaryocytes. Dot size represents the ratio of genes associated with each pathway, while color intensity indicates statistical significance, expressed as -log10 adjusted p-value. (G) Representative fluorescence images of cells cultured in a 2D plate and 3D SilkInk. Cells were stained for CD61 and then shown using a pseudocolor depth scale, where color indicates the z-position of each cell within the imaged volume, expressed in micrometers. 3D SilkInk shows increased formation of MK-forming platelets compared with 2D plate culture. (H) Quantification of MK-forming platelets across the indicated culture conditions (Y-27632 = ROCK inhibitor). (I) Platelet (PLT) output, expressed as fold change relative to 2D plate culture. *p < 0.05; ***p < 0.001; ****p < 0.0001. Data are shown as mean ± SD.

### Megakaryopoiesis in 2D proceeds under chronic genomic stress

Megakaryocyte differentiation occurred within markedly distinct molecular contexts in 2D and SilkInk. Stage-resolved pathway analysis showed that, in 2D, early progenitors, late progenitors, and megakaryocytes all remained enriched for stress programs, TP53-regulated transcription, p53 signaling, G1/S regulation, intrinsic apoptosis, and DNA damage response (**Figure 4A**). Differentiation in 2D, therefore, proceeds under persistent cell-cycle engagement and chronic stress signaling. Direct genomic-instability assays corroborated this: comet assays revealed substantially greater fragmented-DNA migration in 2D than in SilkInk (**Figure 4B**), and γH2AX immunostaining identified a higher fraction of double-strand-break-positive cells in flat culture (**Figure 4C**).

**FIGURE 4.**
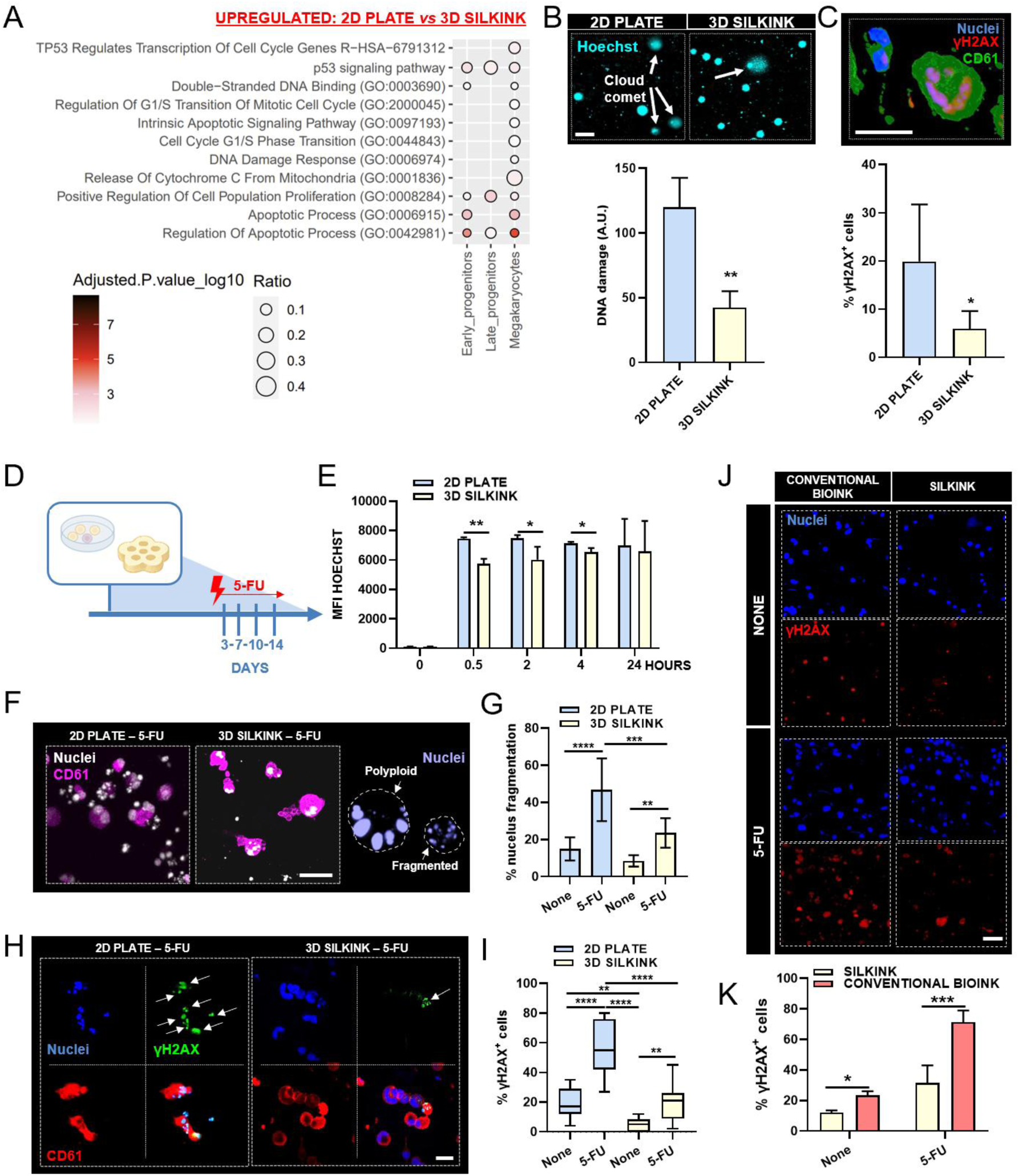
The 3D SilkInk modulates the activation of DNA damage. (A) Ontology/pathway terms enrichment dot plot showing programs upregulated in 2D plates *versus* 3D SilkInk cultures, including p53 signaling, DNA damage responses, apoptosis, and cell-cycle-related pathways, mapped across early progenitors, late progenitors, and megakaryocytes. (B) Representative Hoechst-stained comet assay images from 2D and 3D cultures and corresponding quantification of DNA damage. Scale bar, 10 µm. (C) Representative immunofluorescence image of γH2AX-positive nuclei and quantification of γH2AX-positive cells in 2D and 3D cultures. Scale bar, 30 µm. (D) Experimental scheme for 5-FU treatment after seeding in 2D or 3D cultures. (E) Time course of Hoechst mean fluorescence intensity (MFI) after dye exposure in 2D plates and 3D SilkInk cultures. *p < 0.05, **p < 0.01. (F) Representative images of nuclei/CD61 staining after 5-FU exposure in 2D and 3D cultures, including examples of polyploid and fragmented nuclei. Scale bars, 30 µm. (G) Quantification of nuclear fragmentation in untreated and 5-FU-treated samples cultured in 2D or 3D. **p < 0.01, ***p < 0.001, ****p < 0.0001. (H) Representative γH2AX/CD61/nuclei images in 5-FU-treated 2D and 3D cultures. The arrow indicates γH2AX-positive nuclei. Scale bar, 30 µm. (I) Quantification of γH2AX-positive cells across untreated and 5-FU-treated 2D and 3D conditions. **p < 0.01, ****p < 0.0001. (J) Representative immunofluorescence images of nuclei and γH2AX staining in conventional bioink and SilkInk cultures under untreated conditions or after 5-FU exposure. Scale bar, 30 µm. (K) Quantification of γH2AX-positive cells in conventional bioink and SilkInk cultures under untreated and 5-FU-treated conditions, showing higher basal and 5-FU-induced DNA damage in conventional bioink compared with SilkInk. *p < 0.05, ***p < 0.001. Data are shown as mean ± SD, or median and range.

To test how these distinct baselines shape responses to exogenous genotoxic stress, we exposed cultures to 5-fluorouracil (5-FU), a pyrimidine analog antimetabolite (**Figure 4D**). To rule out diffusion-limited drug delivery as a confounder, we used Hoechst as a small-molecule tracer ^52^. The signal in the 3D matrix rose more gradually but reached levels comparable to 2D within 24 hours (**Figure 4E**). Despite comparable exposure, cellular outcomes diverged sharply. 5-FU produced dose-dependent cell death and impaired platelet output in 2D, whereas SilkInk-encapsulated HSPCs remained significantly more resistant (**Supplemental Figure 5**). We therefore used 0.75 μM 5-FU, a physiologically relevant antimetabolite stress, in subsequent experiments.

Morphological analysis reinforced these differences. After treatment, 2D cultures showed frequent nuclear fragmentation, while SilkInk preserved large polyploid megakaryocytes (**Figures 4F and 4G**). γH2AX staining in CD61^+^ megakaryocytic cells confirmed that 5-FU sharply amplified DNA damage signaling in 2D cultures but was significantly attenuated in SilkInk (**Figures 4H and 4I**). Critically, this protection was not a generic consequence of 3D culture: a conventional hydrogel bioink showed higher basal γH2AX positivity than SilkInk and accumulated substantially greater 5-FU-induced damage (**Figures 4J and 4K**). Generic 3D matrices were thus insufficient to suppress the stress phenotype; SilkInk’s specific biomechanical and biochemical properties were required.

The same hierarchy held for differentiation. Ploidy analysis and 3D volumetric reconstruction showed that SilkInk supported progression toward larger cell volumes and higher ploidy, with increased frequencies of 8N and >16N megakaryocytes, whereas the conventional bioink retained predominantly small, low-ploidy cells (**Supplemental Figures 6A-C**). High-resolution imaging confirmed increased proplatelet formation, branching complexity, and CD61^+^ platelet-sized particle accumulation in SilkInk relative to the conventional bioink (**Supplemental Figures 6D-F**). Compared with the conventional bioink used as a reference, which displays a viscosity of 2.6-7.5 kPa·s at 0.01 s^−^¹, SilkInk showed a much lower low-shear viscosity, approximately 8-10 Pa·s at 0.01 s^−^¹ (**Supplemental Figure 1**), corresponding to an approximate maximal 900-fold reduction in viscosity. At high shear rate, the two materials were more comparable, with SilkInk reaching approximately 1 Pa·s and the conventional bioink reported at 1.0-1.9 Pa·s at 200 s^−^¹, indicating that SilkInk combines shear-thinning behavior with an exceptionally soft and low-viscosity environment that sustains the endomitotic growth program required for productive megakaryopoiesis.

### SilkInk resolves stress-impaired thrombopoiesis as a recoverable functional state

The relationship between DNA-damage signaling and thrombopoiesis is context-dependent, shaped by the magnitude, persistence, and resolution of stress ^4,6^. Controlled DDR activation may contribute to megakaryocyte maturation, whereas excessive or unresolved damage impairs thrombopoiesis ^4^. Because 2D culture and non-silk 3D matrices impose elevated basal stress, persistent DNA-damage signaling, and altered cell-cycle regulation on megakaryopoiesis, the baseline state likely shapes not only the magnitude of antimetabolite injury but also the system’s capacity to recover after drug withdrawal. To test this, we exposed cultures to 5-FU followed by washout. In 2D, 5-FU rapidly drove cells beyond a reparable threshold, with marked loss of viability and near-complete suppression of polyploid megakaryocyte generation (**Figure 5A**); drug removal failed to restore either viability or polyploidization, indicating that 2D culture converts transient antimetabolite exposure into sustained cytotoxic collapse. In SilkInk, the response was graded: 5-FU reduced viability and polyploid output, but washout produced substantial recovery of both, with cultures progressively approaching untreated baselines (**Figure 5B**). SilkInk therefore preserves a recoverable thrombopoietic state that 2D culture obscures through excessive culture-induced injury. This *in vitro* behavior matched the *in vivo* response to transient 5-FU. In mice, treatment induced a pronounced but reversible reduction in marrow cellularity with progressive recovery at later time points (**Figures 5C and 5D**). Peripheral platelet counts declined and rebounded (**Figure 5E**), the proportion of polyploid megakaryocytes was preserved throughout (**Figure 5F**), and histological analysis confirmed megakaryocyte persistence in the marrow during both injury and recovery phases (**Figure 5G**). 5-FU, therefore, transiently disrupts thrombopoietic function without ablating the megakaryocytic lineage, exactly the pattern SilkInk recapitulates and 2D does not. SilkInk also enabled direct visualization of proplatelet formation, a highly dynamic process that is difficult to quantify *in vivo*. Live imaging showed that SilkInk-resident megakaryocytes undergo progressive cytoplasmic remodeling and proplatelet extension (**Figure 5H**). 5-FU markedly impaired proplatelet formation, and washout significantly restored it (**Figures 5I and 5J**). Platelet output followed the same trajectory, declining after 5-FU and partially recovering after washout (**Figure 5K**).

**FIGURE 5.**
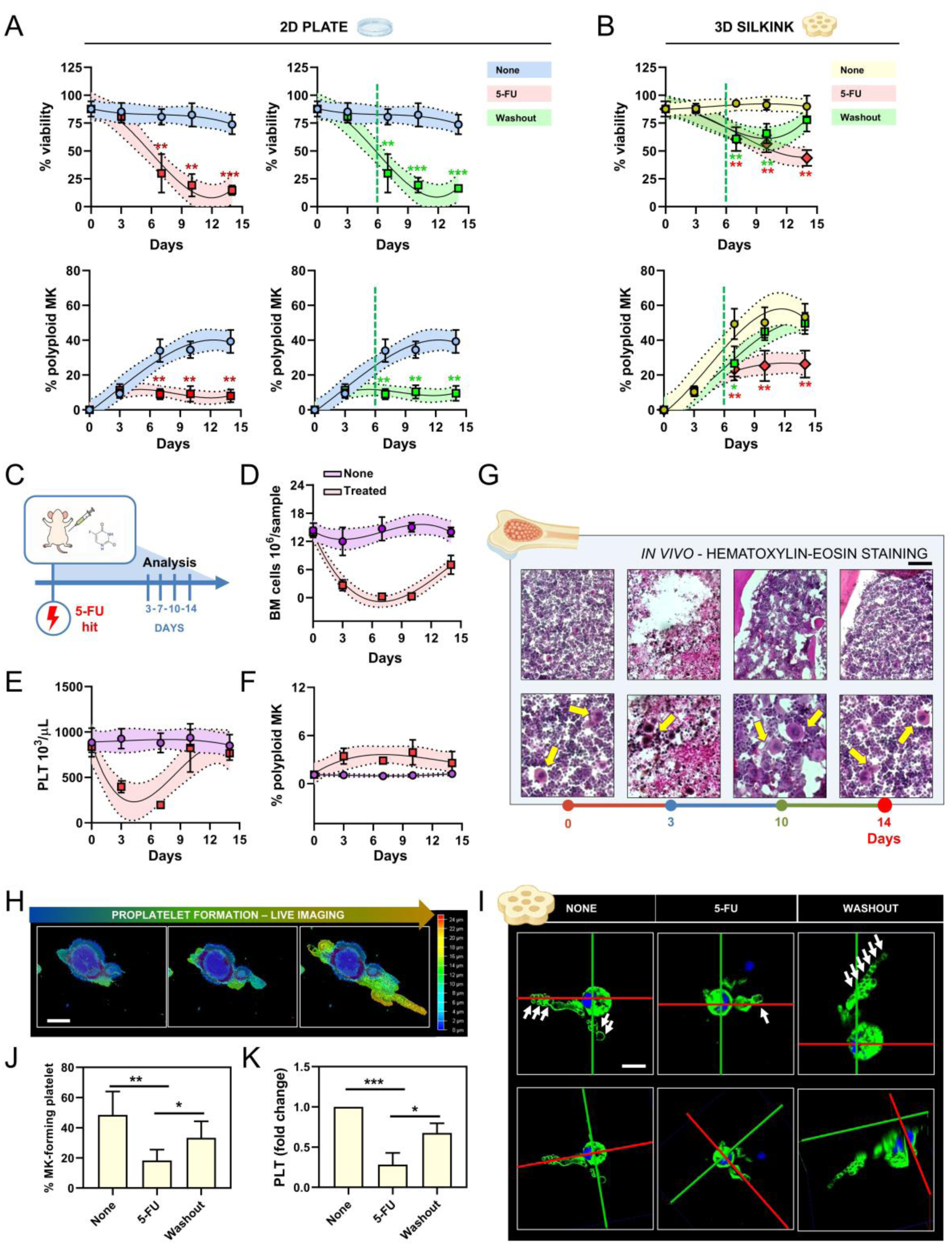
3D SilkInk captures reversible impairment of thrombopoiesis after antimetabolite stress. (A) Time-course analysis of viability and polyploid megakaryocyte generation in 2D plate cultures under untreated conditions (blue), continuous 5-FU exposure (red), or 5-FU washout (green). The dashed green line indicates the time at which the drug is removed. **p < 0.01, ***p < 0.001. (B) Time-course analysis of viability and polyploid megakaryocyte generation in 3D SilkInk cultures under untreated conditions (yellow), continuous 5-FU exposure (red), or 5-FU washout (green). The dashed green line indicates the time at which the drug is removed. *p < 0.05, **p < 0.01. (C) *In vivo* experimental scheme for 5-FU administration and longitudinal analysis at the indicated time points. (D) Quantification of bone marrow cellularity over time after 5-FU treatment compared with untreated controls. (E) Quantification of peripheral platelet counts over time after 5-FU treatment compared with untreated controls. (F) Frequency of polyploid megakaryocytes in bone marrow over time after 5-FU treatment. (G) Representative hematoxylin and eosin-stained bone marrow sections at the indicated time points after 5-FU exposure, with megakaryocytes indicated by arrows. Scale bar, 200 µm. (H) Representative live-imaging sequence showing progressive cytoplasmic remodeling and proplatelet extension by a megakaryocyte in 3D SilkInk. The megakaryocyte was stained for CD61 and then shown using a pseudocolor depth scale, where color indicates the z-position of each part of the cell body/branch of the imaged volume, expressed in micrometers. Scale bar 50 µm. (I) Representative 3D confocal reconstructions of megakaryocytes in 3D SilkInk under untreated, 5-FU-treated, and washout conditions, showing suppression and recovery of proplatelet-forming structures. Scale bar, 30 µm. (J) Quantification of proplatelet-forming megakaryocytes in 3D SilkInk under untreated, 5-FU-treated, and washout conditions. (K) Platelet output from 3D SilkInk cultures under untreated, 5-FU-treated, and washout conditions, expressed as fold change relative to untreated controls. *p < 0.05, **p < 0.01, ***p < 0.001. Data are shown as mean ± SD.

Together, these data show that SilkInk resolves stress-impaired thrombopoiesis as a reversible functional state rather than a nonspecific cytotoxic endpoint, providing a reproducible model for studying platelet production, marrow recovery, and drug-induced hematopoietic injury.

### TPO-receptor agonists pharmacologically rescue stress-impaired thrombopoiesis in SilkInk

Accumulating evidence identifies the TPO signaling axis as a stress-adaptive pathway in hematopoietic progenitors. TPO actively promotes DNA repair in HSPCs via DNA-PK-dependent non-homologous end joining (NHEJ), limiting long-term genotoxic injury ^53^. The TPO-receptor agonist (TPO-RA) eltrombopag enhances double-strand break repair through canonical NHEJ, improving genome stability and survival of human HSPCs ^54^; romiplostim confers hematopoietic protection in severe genotoxic settings, including lethal irradiation, consistent with a broader capacity to support hematopoietic recovery under DNA-damaging stress ^55,56^.

We therefore tested whether the SilkInk niche could support pharmacological rescue of antimetabolite-impaired thrombopoiesis. SilkInk retained an active TPO-responsive signaling axis: both romiplostim and eltrombopag increased AKT and ERK phosphorylation and shifted the apoptotic balance toward survival, with elevated BCL-XL relative to BAK and BAX (**Figures 6A-6E**). To target the developmental window most vulnerable to 5-FU, TPO-RAs were administered during the late-progenitor-to-megakaryocyte transition (**Figure 6F**). Both agents preserved a survival-biased BCL-XL/BAK and BCL-XL/BAX balance under 5-FU challenge, counteracting drug-induced apoptotic pressure (**Figures 6G-6I**). This protection was accompanied by reduced genomic stress. Romiplostim and eltrombopag reduced baseline γH2AX positivity in CD61^+^ cells and attenuated γH2AX accumulation after 5-FU (**Figures 6J-M**), indicating dampened DNA damage signaling during megakaryocyte differentiation. The molecular rescue translated into functional recovery: relative to 5-FU alone, both TPO-RAs improved viability, restored polyploid megakaryocyte formation, and enhanced proplatelet-forming capacity (**Figures 6N-P**). SilkInk therefore captures stress-impaired thrombopoiesis as a reversible, pharmacologically responsive trajectory spanning injury, adaptation, and recovery.

**FIGURE 6.**
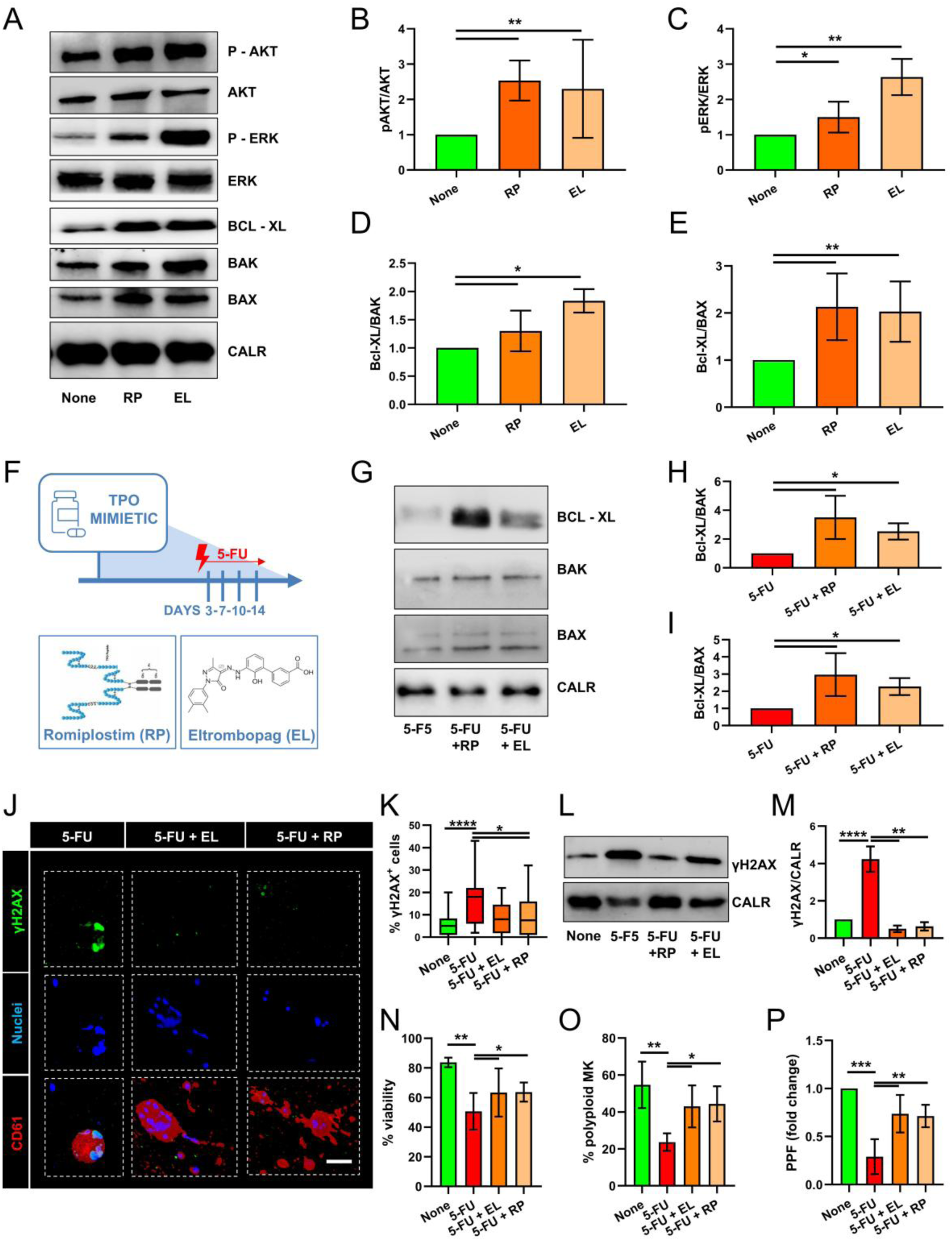
3D SilkInk captures TPO-receptor agonist-mediated rescue of stress-impaired thrombopoiesis. (A) Representative immunoblot analysis of AKT, phospho-AKT, ERK, phospho-ERK, BCL-XL, BAK, and BAX in samples treated with romiplostim (RP) or eltrombopag (EL), with calreticulin (CALR) as loading control. (B-C) Densitometric quantification of phospho-AKT/AKT and phospho-ERK/ERK ratios, showing activation of canonical TPO-responsive signaling after RP or EL treatment. *p < 0.05, **p < 0.01. (D-E) Quantification of BCL-XL/BAK and BCL-XL/BAX ratios, indicating a shift toward pro-survival apoptotic control after TPO-receptor agonist treatment. *p < 0.05, **p < 0.01. (F) Experimental scheme for TPO-receptor agonist treatment during 5-FU challenge in SilkInk cultures. (G) Representative immunoblot analysis of BCL-XL, BAK, and BAX in 5-FU-treated cultures with or without RP or EL, with calreticulin as loading control. (H-I) Quantification of BCL-XL/BAK and BCL-XL/BAX ratios after 5-FU exposure, showing preservation of a survival-biased apoptotic balance upon RP or EL treatment. *p < 0.05. (J) Representative immunofluorescence images of γH2AX, nuclei, and CD61 in SilkInk cultures treated with 5-FU alone or in combination with EL or RP. Scale bar, 30 µm. (K) Quantification of γH2AX-positive cells across untreated, 5-FU, 5-FU + EL, and 5-FU + RP conditions, showing reduced DNA damage upon TPO-receptor agonist treatment. (L) Representative immunoblot analysis of γH2AX in untreated, 5-FU, 5-FU + RP, and 5-FU + EL conditions, with calreticulin as loading control. (M) Densitometric quantification of γH2AX/calreticulin ratio, confirming attenuation of 5-FU-induced DNA damage after RP or EL treatment. *p < 0.05, **p < 0.01; ****p < 0.0001. (N-P) Functional assessment of thrombopoietic recovery in SilkInk cultures, showing viability, polyploid megakaryocyte frequency, and proplatelet formation across untreated, 5-FU, 5-FU + EL, and 5-FU + RP conditions. *p < 0.05, **p < 0.01, ***p < 0.001. Data are shown as mean ± SD, or median and range.

## Discussion

Hematopoietic cells reside in a mechanically distinctive niche, but most *in vitro* studies remove them from it. Our findings show that this choice has not been biologically neutral: conventional 2D culture imposes a chronic, mechanotransduction-driven state on HSPCs, characterized by sustained replication stress, persistent genome-surveillance signaling, and redox imbalance, that distorts hematopoietic biology and likely accounts for a substantial share of what has been interpreted as intrinsic *in vitro* stress ^1,2,14,57–59^.

By providing the viscoelastic compliance and platelet-inert chemistry of a marrow-like microenvironment, SilkInk reveals that niche mechanics are an active determinant of how hematopoietic cells engage stress signaling and execute differentiation. This conclusion has direct implications for ongoing efforts to develop human-relevant, ethical, and mechanistically informative models of hematopoiesis ^60–62^. Organoid, organ-on-chip, and biomaterial-based platforms have substantially advanced bone marrow engineering ^63–65^, yet sustaining functionally competent hematopoiesis *in vitro* has remained difficult, with most systems constrained by reproducibility, scalability, or dependence on heterogeneous stromal support.

SilkInk addresses these constraints with a chemically defined, mechanically tunable silk-fibroin formulation that recapitulates marrow viscoelasticity, supports printable and reproducible 3D constructs, and, critically for hematopoietic applications, does not activate platelets. Within this framework, megakaryopoiesis emerged as a particularly informative readout, because lineage maturation depends on coordinated cytoskeletal remodeling, endomitosis, and platelet biogenesis within compliant perisinusoidal niches ^3,27,65–71^, and is therefore acutely sensitive to substrate mechanics. Our data support a threshold model in which the biological consequences of DNA-damage signaling depend less on its presence than on the mechanical and metabolic context in which it operates ^10,76–78^. In SilkInk, DDR activation remained compatible with differentiation, allowing megakaryocytes to progress through endomitosis and proplatelet formation while preserving adaptive function. In 2D, sustained mechanotransduction accompanied chronic replication-stress signaling, persistent DSB-repair activation, and elevated baseline γH2AX, a background against which DNA-damage response crossed the threshold from adaptive to pathological ^72–74^.

Critically, a conventional non-silk hydrogel also failed to protect against this transition, indicating that 3D dimensionality alone is insufficient; the specific viscoelastic and platelet-inert properties of silk are required to decouple genome surveillance from genotoxic collapse. Niche mechanics, therefore, act not as a permissive backdrop but as an active regulator of how hematopoietic cells calibrate stress against differentiation. The threshold effect became most evident under antimetabolite challenge. 5-fluorouracil has long served as a model of stress hematopoiesis because it preferentially targets cycling progenitors and elicits a regenerative phase after injury ^43,75–79^. In our system, the response was fully determined by the baseline culture state: 2D cells, already operating under elevated chronic stress, entered sustained cytotoxic collapse with loss of polyploidization, suppression of proplatelet formation, and irreversible loss of platelet output even after drug withdrawal. SilkInk-encapsulated cultures, by contrast, recovered after washout, recapitulating the transient, regenerative pattern observed *in vivo*, where marrow cellularity, peripheral platelet counts, and polyploid megakaryocyte populations declined and then rebounded. This parallel between SilkInk and *in vivo* biology, and its absence in 2D, is what makes the *in vitro* system useful for interrogating recovery rather than injury alone. The response to TPO-receptor agonists tested this directly. TPO signaling has been linked to both megakaryocyte maturation and genome maintenance in HSPCs, and recent work shows that TPO and TPO-RAs enhance DNA repair, reduce γH2AX-associated injury, and improve survival under genotoxic stress ^53–56^. Within SilkInk, both eltrombopag and romiplostim engaged active AKT and ERK signaling, shifted the apoptotic balance toward survival through elevated BCL-XL relative to BAK and BAX, reduced 5-FU-induced DNA damage in CD61^+^ cells, restored polyploid megakaryocyte formation, and enhanced proplatelet output. The value of these rescues lies not in improving isolated readouts but in demonstrating that SilkInk preserves an injury-adaptation-recovery trajectory in which mechanism, not just outcome, can be dissected.

SilkInk combines several features that extend its utility across multiple settings of perturbed human megakaryopoiesis. The platform resolves substrate-tunable HSPC stress signaling, captures endomitotic and thrombopoietic trajectories under genotoxic challenge, and supports pharmacologically responsive dynamics of injury and recovery. Relevant applications include inherited thrombocytopenia and bone marrow failure syndromes linked to DDR or replication-stress pathways ^4–6^, myeloproliferative neoplasms with thrombocytopenic phenotypes, and radiation-induced hematopoietic injury. The system also offers value in preclinical screening of myelosuppressive drug candidates, where interspecies differences in megakaryopoiesis have repeatedly limited the predictive value of murine models. Chemotherapy-induced thrombocytopenia (CIT) is a particularly well-defined application ^80^. Clinical management remains inconsistent, and patients face competing risks from cancer-associated hypercoagulability and bleeding from myelosuppressive treatment ^81^. Open questions about how emerging antineoplastics, such as PARP inhibitors, interact with TPO-RA support are clinically pressing yet mechanistically underexplored ^82,83^. By preserving human HSPC self-renewal, lineage-specific differentiation, and pharmacological responsiveness, SilkInk provides a substrate for addressing these questions.

A few limitations frame these conclusions. SilkInk in its current configuration lacks the cellular complexity of the native marrow, mesenchymal stromal, osteoblastic, and endothelial cell components, as well as sinusoidal architecture, and does not incorporate active perfusion or oxygen gradients. The translational arguments rest on a single antimetabolite (e.g., 5-FU), and generalization to other myelosuppressive classes will need direct testing. These caveats define the next phase of the work rather than undermine the present conclusions.

Taken together, our findings establish niche mechanics as an active determinant of whether hematopoietic stress remains adaptive or becomes pathological and identify SilkInk as a reproducible system in which the biology of human marrow injury and recovery can be resolved at mechanistic depth. SilkInk enables what neither 2D culture nor animal models allow. It provides simultaneous, mechanistic resolution of injury, adaptation, and recovery in a human hematopoietic system that respects the physical biology of the niche.

## Acknowledgments

This paper was supported by the EIC Transition Project HORIZON-EIC-2022-TRANSITION-01-n. 101113073 SilkInk to AB, and Italian Ministry of University and Research (PRIN 2022-2022P9RM9M) to AB and MMe.

## Author Contributions

C.A.D.B. conducted experiments, acquired and analyzed data, and wrote the manuscript; M.Mi., M.Me., E.P., A.M., S.D., G.G., M.S. conducted experiments, acquired and analyzed data, and edited the manuscript; C.D.F. provided blood samples; V.K. provided technical support, analyzed data, and edited the manuscript; M.Ma. provided scientific and technical support; A.B. conceived the idea, supervised the project, designed research studies, acquired and analyzed data, and wrote the manuscript.

## Conflict of Interest

The authors declare no conflict of interest.

## Notes

### Competing Interest Statement

The authors have declared no competing interest.

