## Supplemental File for "A bioprinted silk marrow niche reveals mechanical regulation of human megakaryopoiesis under genotoxic stress"

### Supplemental Materials and Methods

***Preparation of silk fibroin bioink and bioprinting parameters.*** Silk fibroin aqueous solution was obtained from *Bombyx mori,* and SilkInk was prepared, as previously described [1,2]. SilkInk was loaded into a sterile cartridge fitted with a 20G nozzle and kept at 37 °C in a BIO-X extrusion bioprinter (Cellink Bioprinting). The thermo-regulated printhead was equipped with an insulator that can house a cylindrical nozzle/needle. This allowed precise control of the temperature over the needle, ensuring uniform temperature down to the print surface while maintaining cells in their physiological condition. The printing conditions were as previously described [2]. We designed the desired pattern in Autodesk Fusion 360. The construct consisted of a layered hexagon intersected with smaller hexagons at its borders to shape a '*flower*'-like model. The use of small grids is intended to enable homogeneous diffusion of the medium within the scaffolds during cell culture. After bioprinting, the scaffold was crosslinked in a physiological salt solution containing $CaCl_2$. Then, the 3D construct was immersed in cell culture medium and kept at 37 °C and 5% $CO_2$.

***Rheology.*** The rheological characterization of SilkInk was performed using the Discovery HR-10 rheometer (TA Instruments) with a bottom Peltier plate and an upper 20 mm Peltier plate geometry with a 500 µm gap. Rheological measurements were performed as technical triplicates on at least three different independent batches, preconditioned at 37 °C before loading and equilibrated in the rheometer for 60 s before initiating measurements:

- Steady-state flow sweeps were performed at 37 °C over a shear rate range of 0.01-500 1/s.

- Thixotropy was tested by subjecting the samples to intervals of low shear (60 s 0.01 1/s) and high shear (10 s 500 1/s) to mimic bioink deformation under pressure during printing and the recovery after the deformation.

- Temperature sweeps were performed between 37 °C and 20 °C and then back up to 37 °C, with a linear ramp (3°C/min) up and keeping the strain (0.1%) and frequency (0.5 Hz) constant.

The characterizations of mechanical properties of crosslinked bioink were performed at 37 °C on the rheometer, on disc-shaped samples with 8 mm diameter and ~0.7-1.4 mm height, bioprinted, ionically crosslinked, and then maintained in cell culture conditions until testing. Four technical replicates were tested for three batches.

- Uniaxial unconfined compression testing was performed using the 20 mm upper plate geometry. The gap was reduced until a positive axial force was detected, and displacement was applied at a rate of 1% strain/s based on the sample height, up to 60% strain, to generate force-displacement curves. These were converted to stress-strain curves using measured sample diameters, and tangent modulus values were extracted via linear regression over the 5-10% strain range.

- Frequency sweeps were carried out with an 8 mm serrated upper geometry to avoid slippage, and the gap was decreased until a 0.005 N force was detected. Oscillation frequency sweeps were performed at 0.1% strain between 0.1 and 1 Hz.

Analyses demonstrated that SilkInk behaves as a shear-thinning fluid (**Supplemental Figure 1A**). Flow sweep analysis revealed that viscosity decreases progressively with increasing shear rate, a profile compatible with extrusion-based bioprinting approaches and that limits excessive shear stress during deposition. Cyclic high-shear challenges indicated thixotropic recovery, with viscosity dropping under high shear and recovering when shear was reduced, supporting the capacity to maintain shape fidelity during extrusion (**Supplemental Figure 1B**). Oscillation temperature sweeps further showed a clear viscoelastic transition. Both storage (G') and loss (G'') moduli declined with increasing temperature and converged around ~28°C, consistent with a formulation that can be easily printed/handled at physiological temperature and rapidly transitions to a stable gel on the cooled print bed, preserving print fidelity (**Supplemental Figure 1C**).

Mechanical reproducibility and temporal stability of the bioprinted SilkInk constructs were assessed using small-amplitude oscillatory shear and uniaxial compression. Oscillation rheology frequency-sweeps showed that 3D-printed constructs behaved as viscoelastic compositions, with the storage modulus (G') increasing monotonically over the 0.1-1 Hz range across three independent batches, indicating a consistent frequency-dependent response and supporting print-to-print reproducibility (**Supplemental Figure 1D**). Across all batches, G' was higher at day 1 and decreased by day 7, revealing a moderate time-dependent softening in shear, attributed to equilibration of $Ca^{2+}$ ions from the ionically crosslinked alginate component with culture media, and melting of the gelatin component at 37 °C. The overall rheological profile was preserved (**Supplemental Figure 1D**). In parallel, compression testing of silk-based constructs revealed non-linear stress-strain behavior with progressive stiffening, a characteristic response of polymeric hydrogel networks, and closely overlapping curves across batches at both time points, consistent with preserved macroscopic integrity after 3D printing, crosslinking, and culture (**Supplemental Figure 1E**).

***Imaging.*** For immunofluorescence imaging, samples were fixed in 4% formaldehyde for 20 min at room temperature. Samples were probed with anti-CD34 (1:100), anti-CD61 (1:100), and anti-γH2AX (Abcam) overnight at 4 °C. Alexa Fluor secondary antibody (1:500) (Invitrogen) was incubated for 2 h at room temperature. Nuclei were stained with Hoechst (Sigma-Aldrich). Samples were imaged by an SP8 confocal laser scanning microscope (Leica). Isotype controls were used as negative controls to exclude nonspecific background signals. The acquisition parameters were set on the negative controls. 3D reconstruction and image processing were performed using Leica licensed software.

***Flow cytometry.*** Megakaryocytes and platelets were retrieved from the silk bioink by using the solution for dissolving the 3D construct described above. Flow cytometry settings were established, as previously described [3]. Megakaryocyte ploidy, surface marker expression, and viability were analyzed, as previously described [4]. Platelets were analyzed using the same forward- and side-scatter patterns as in human peripheral blood platelets. Isotype controls were used as negative controls to exclude nonspecific background signals. DNA synthesis and S-phase entry in megakaryocytes were assessed using the EdU Assay/EdU Staining Kit (iFluor 488, ab219801, Abcam) according to the manufacturer's instructions to identify EdU-incorporating polyploid megakaryocytes. All samples were acquired with a BD FACS Lyric (Becton Dickinson) flow cytometer. Off-line data analysis was performed using Kaluza software package (Beckman Coulter).

***Animals.*** 6-8-week-old C57/BL6 mice were obtained from Charles River Laboratories, Italy. Mice were housed at the animal facility of the Department of Physiology, section of General Physiology, University of Pavia (approval #1/2010, 24/06/2010). All animals were sacrificed in accordance with current European legal requirements for animal practice. Myelosuppression was induced by 5-fluorouracil (5-FU) (250 mg/kg body weight, Sigma-Aldrich, Milan, Italy) injected intraperitoneally. Age-paired mice were injected with PBS as a control. At the indicated point, mice were sacrificed, and bones and blood were collected for analyses. Femurs were fixed for 24 hours in 3% paraformaldehyde (PFA) and decalcified in 10% EDTA in phosphate-buffered saline (PBS) (w/o calcium and magnesium) at pH 7.2 for 2 weeks at 4°C. Specimens were embedded in optimal cutting temperature (OCT) cryosectioning medium and snap frozen in a chilling bath. Eight-micrometer tissue sections were prepared using a Microm Microtome HM 250 (Bio-Optica S.p.A., Milan, Italy, www.bio-optica.it).

***Western Blot Analysis.*** Samples were lysed with HEPES-glycerol lysis buffer (50 mM HEPES, 150 mM NaCl, 10% glycerol, 1% Triton X-100, 1.5 mM $MgCl_2$,1 mM EGTA, 10 mM NaF, 1 mM $Na_3VO_4$, 1 mg/mL leupeptin, 1 mg/mL aprotinin). Lysis was performed on ice for 30 min, and lysates were clarified by centrifugation at 14,000 × g for 15 min at 4 °C. Protein lysates were subjected to sodium dodecyl sulfate–polyacrylamide gel electrophoresis in 12% acrylamide gels and transferred to polyvinylidene fluoride membrane (Bio-Rad). Membranes were probed with affinity-purified primary antibodies against phospho-AKT, AKT, phospho-ERK, ERK, Bcl-Xl, Bak, and Bax (1:1000; Life Technologies) [5]. Anti-calreticulin antibody (1:1000, Abcam) was used to ensure equal loading. Immunoreactive bands were detected by horseradish peroxidase-labeled secondary antibodies using enhanced chemiluminescence reagent (Merck-Millipore). Pre-stained protein ladders were used to estimate the molecular weights of the protein of interest (BioRad).

***Comet Assay to measure the DNA damage.*** DNA damage was assessed using the comet assay. After washing with PBS, the cells were embedded in 1% (w/v) low-melting-point agarose (Gibco) and immediately transferred onto glass microscope slides precoated with 1% (w/v) standard agarose (Gibco). The cells were then lysed in a solution containing 2.5 mM NaCl, 0.1M EDTA, 10 mM Tris-base, 1% Triton X-100 (pH 10), for 2h at 4° C. Thereafter, slides were equilibrated with electrophoresis buffer (0.3 M NaOH, 1 mM EDTA) for 40 min to unwind the DNA and then electrophoresed for 30 min at 25V. The nucleoids were subsequently washed in neutralizing buffer (0.4 M Tris-HCl, pH 7.5) and stained with Hoechst 33258 (5µg/ml) (Collins 2004). Hoechst-stained nucleoids were visualized using a fluorescence microscope (Nikon Eclipse E400) with a 40x magnification or a SP8 confocal laser scanning microscope (Leica). Samples were analyzed in a blind manner. For each slide, 100 nucleoids were scored and classified into arbitrary units based on the grade of damage, according to previously published literature [6].

***Proteomics Sample Preparation and LC-MS/MS Analysis.*** Cells from 2D cultures or SilkInk were homogenized using the EasyPep MS Sample Kit (Thermo Scientific Pierce). Following lysis and digestion, the resulting peptides were resuspended in 0.1% formic acid. Peptide concentration was determined via a quantitative colorimetric peptide assay (Thermo Fisher Scientific). For the proteomic run, 2 µg of each peptide sample was loaded onto an UltiMate 3000 RSLC nano system integrated with an Exploris 240 mass spectrometer (Thermo Fisher Scientific). Chromatographic separation was achieved using an Easy-Spray Pepmap RSLC C18 column (2um x 50cm x 75um) at a constant flow rate of 250 nL/min. The mobile phase gradient (Solvent B: 80% acetonitrile/0.1% formic acid; Solvent A: 0.1% formic acid in water) transitioned from 2% to 40% over 150 minutes, followed by a ramp to 99% B in 20 minutes. After a 10-minute hold at 99% B, the column was re-equilibrated for 10 minutes.

Mass spectrometry data were collected in positive ion mode using a data-dependent acquisition (DDA) strategy. Full MS1 scans were performed across an m/z range of 350–1500 at a resolution of 120,000 (at m/z 200), with an AGC target of 3e6 and automatic injection timing. MS2 fragmentation was triggered for precursor ions with intensities exceeding 1e4. Higher-energy collisional dissociation (HCD) was employed with a normalized collision energy of 30%, an AGC target of 7.5e4, and a 40ms maximum injection time. MS2 resolution was set to 15,000, and internal calibration was applied at the start of each run. Each biological sample was processed in five technical replicates.

***Bioinformatic and Data Processing.*** Raw sequencing data were demultiplexed using the mkfastq function from Cell Ranger (v6.1.2, 10x Genomics). Sequencing reads were aligned to the human reference genome (GRCh38; 10x Genomics reference version 2020-A) using the Cell Ranger count pipeline [7].

The resulting gene expression matrices were imported into R and processed using the Seurat package (v5.0.0) [8]. Unless otherwise specified, default parameters were applied. Cells were retained if they expressed at least 200 genes, based on genes detected in at least 3 cells across the dataset. Cells with more than 25% of reads mapping to mitochondrial genes were excluded.

After quality-control filtering, a total of 17,650 single cells were retained, including 11,793 from 2D samples and 5,857 from 3D samples. The datasets were normalized and scaled using the standard Seurat workflow, then integrated with Harmony (v0.1.1) [9].

Dimensionality reduction was performed using principal component analysis (PCA), and the number of principal components was selected based on inspection of the ElbowPlot (n = 50 PCs). Following integration, UMAP was applied for visualization using the RunUMAP function (dims = 1:30). Cell clustering was carried out using the FindNeighbors and FindClusters functions with a resolution of 0.1. Clusters were annotated based on the expression of canonical marker genes and visualized using the DotPlot function.

Cluster-specific marker genes and differentially expressed genes (DEGs) between 3D and 2D samples within each cluster were identified using the FindAllMarkers and FindMarkers functions, respectively.

Significant DEGs were defined using a threshold of adjusted p-value $< 0.01$ and $|\log2$ fold change$| > 1$, and were further separated into upregulated and downregulated gene sets.

Gene lists were analyzed using the Enrichr (v3.1) [10] framework across multiple databases, including Gene Ontology (Biological Process, Molecular Function, Cellular Component), KEGG, and

Reactome. Enrichment results were filtered based on an adjusted p-value < 0.05. Selected representative terms were visualized using dot plots generated with ggplot2 (v3.4.4).
Trajectory inference analysis was performed using the Monocle3 (v1.4.26) package [11]. Unless otherwise specified, default parameters were applied. The trajectory graph was learned using the learn_graph function. Cells were ordered along the trajectory using the order_cells function, selecting root nodes corresponding to early progenitor cells based on cluster annotation. Pseudotime values were then assigned to each cell and visualized on the UMAP embedding.
Genes associated with pseudotime progression were identified using the graph_test function based on the principal graph, and significantly associated genes were selected using a threshold of q-value < 0.05 and Moran's I > 0.5. The expression dynamics of selected genes along pseudotime were visualized as a heatmap with the pheatmap package (v1.0.13), using normalized expression values ordered by pseudotime.

***Statistics.*** Values were expressed as mean plus or minus the standard deviation (mean ± SD), or median and range. The Student's t-test or ANOVA, followed by a Bonferroni posttest, was used to analyze the experiments. A value of at least $p < 0.05$ was considered statistically significant.

### Supplemental Figures and Figure Legends

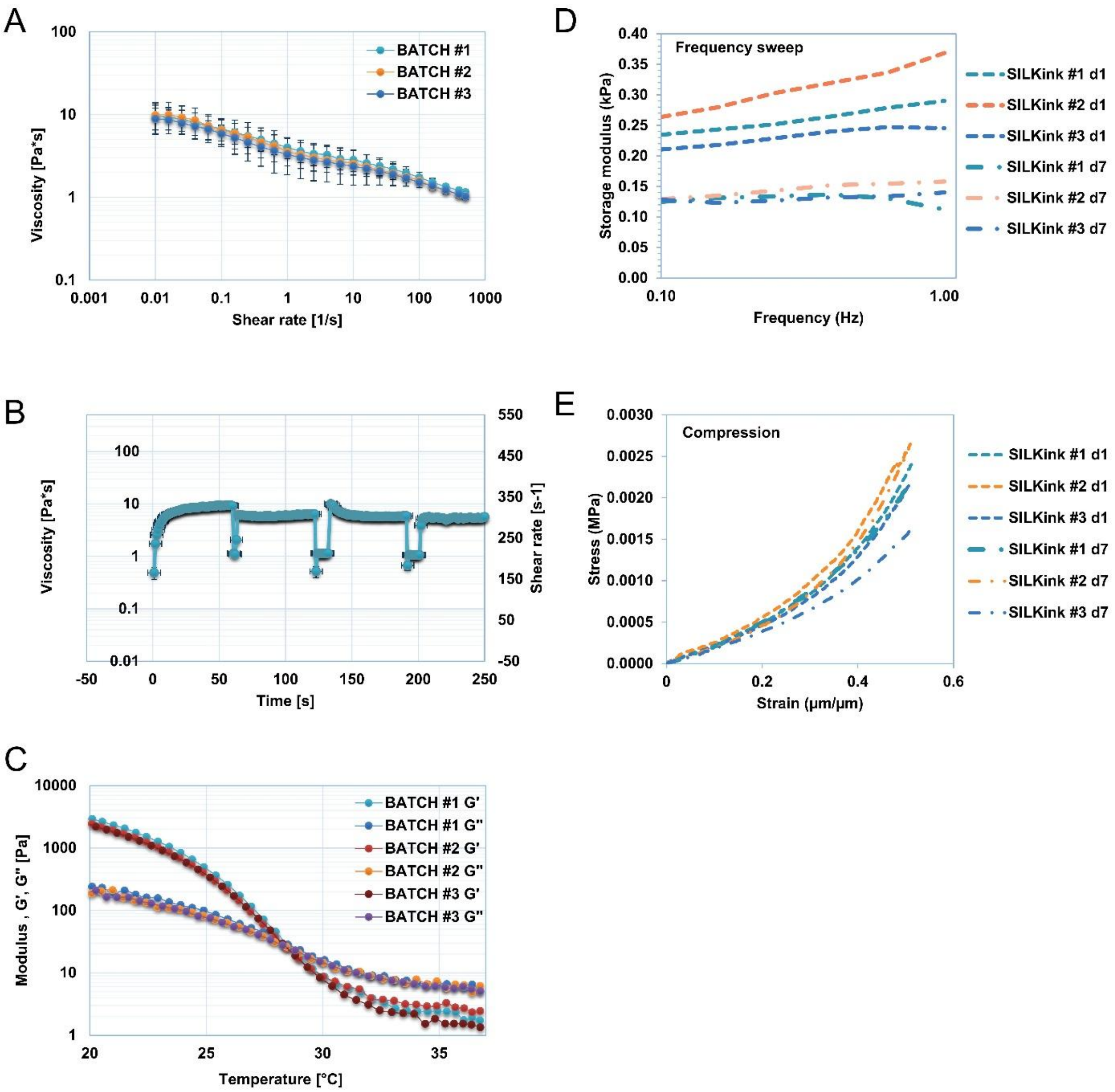


**SUPPLEMENTAL FIGURE 1. Batch-to-batch rheological behavior and mechanical stability of SilkInk formulation and resulting constructs.** (A) Flow curves showing viscosity as a function of shear rate for three independent SilkInk batches. (B) Time-resolved viscosity profile during sequential shear-rate steps, illustrating shear-thinning behavior and recovery. (C) Temperature-sweep reporting storage (G′) and loss (G″) moduli for three independent batches. (D) Frequency sweep showing storage modulus at day 1 and day 7 for three independent constructs. (E) Representative compression stress-strain curves obtained at day 1 and day 7. Data are shown as mean ± SD.

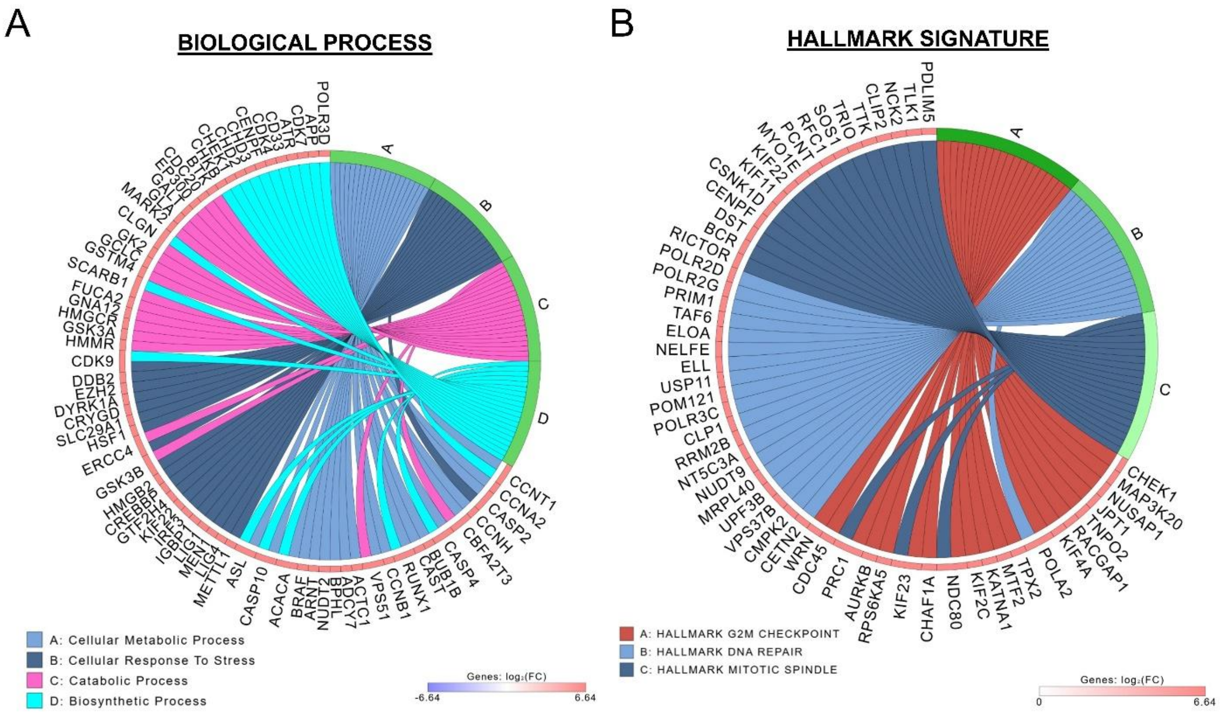


**SUPPLEMENTAL FIGURE 2. Proteomic pathway organization in 2D *versus* 3D cultures.** (A) Chord diagram summarizing enriched biological processes among differentially expressed proteins. (B) Chord diagram summarizing enriched hallmark gene set signatures from the same differential proteomic analysis.

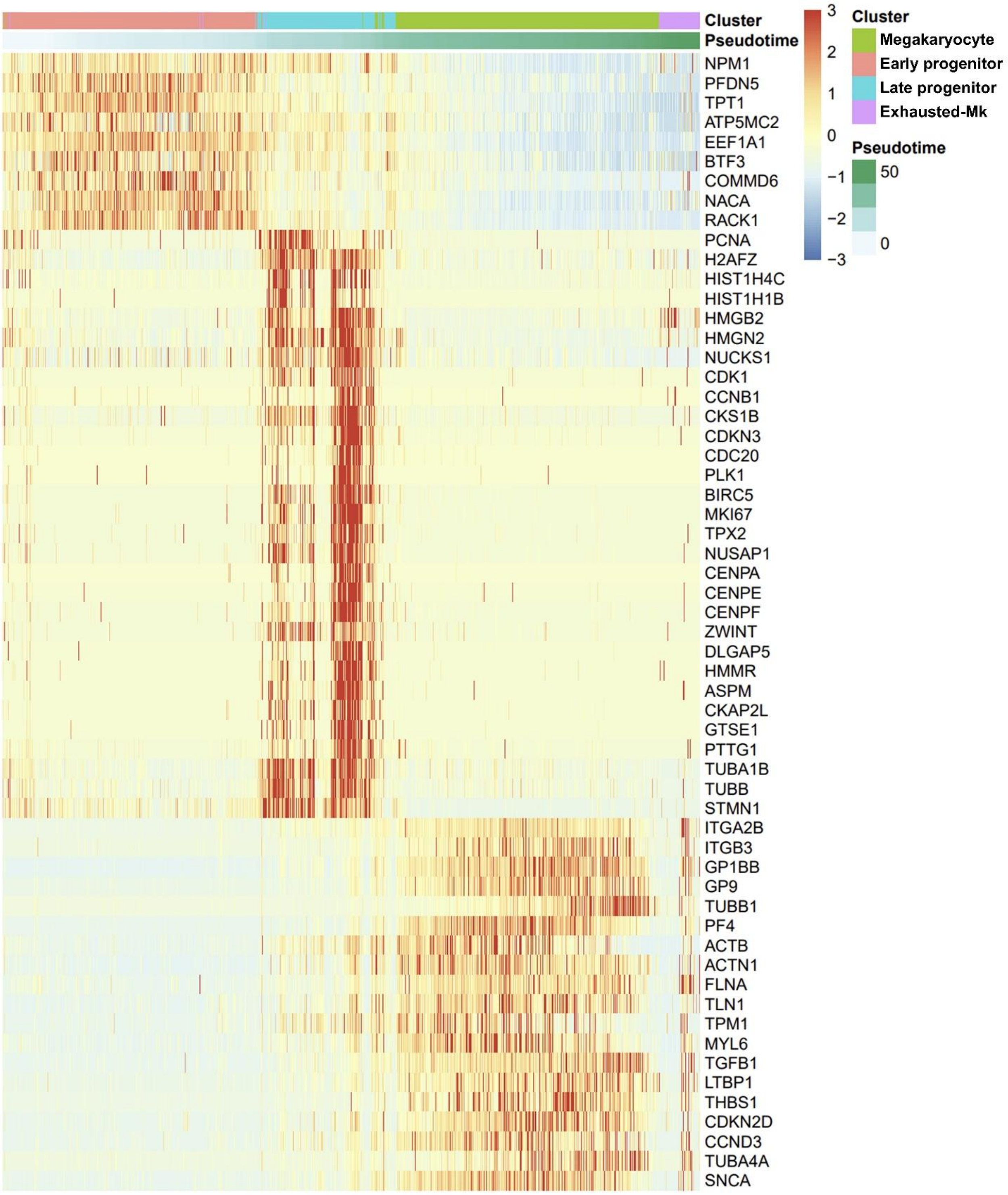


**SUPPLEMENTAL FIGURE 3. Single-cell transcriptional dynamics across megakaryocyte differentiation.** Heat map of representative genes ordered along pseudotime and annotated by cell cluster, showing the transition from early progenitors to late progenitors, megakaryocytes, and exhausted megakaryocytes.

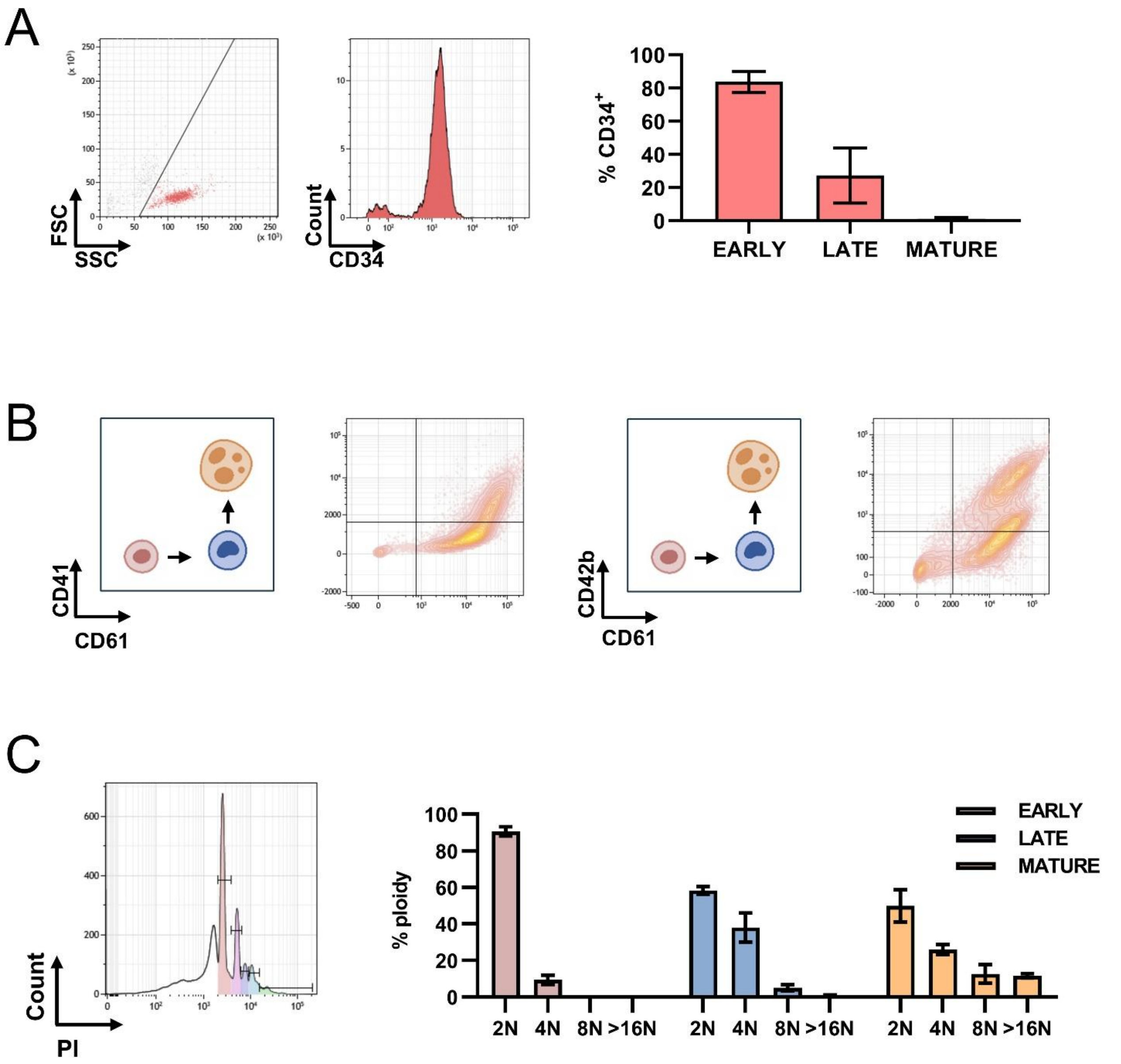


**SUPPLEMENTAL FIGURE 4. Characterization of megakaryocyte differentiation.** (A) Frequency of CD34-positive cells across early progenitors (day 3), late progenitors (day 7), and megakaryocyte fractions (day 10). (B) Representative gating strategy using CD41, CD61, and CD42b to define the indicated differentiation stages. (C) Ploidy distribution (2N, 4N, 8N, and >16N) across early progenitor, late progenitor, and megakaryocyte populations.

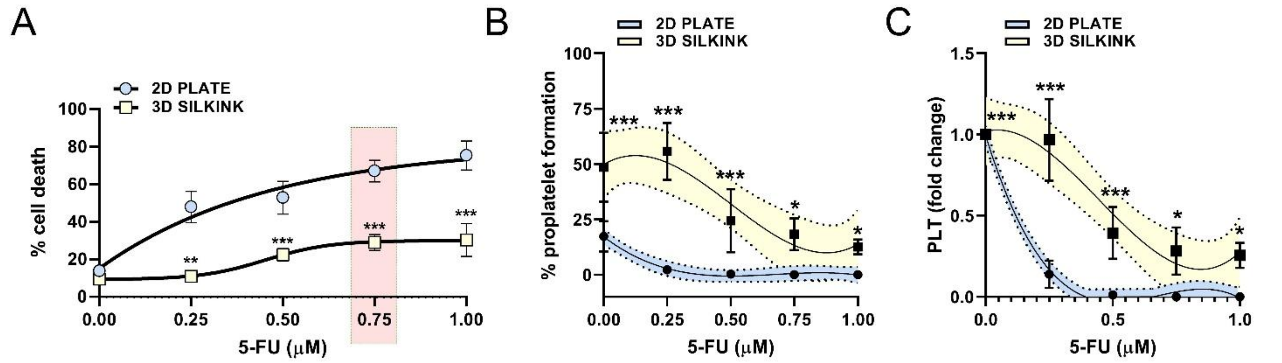


**SUPPLEMENTAL FIGURE 5. 3D SilkInk limits 5-FU-induced cytotoxicity and preserves thrombopoietic output.** (A) Dose-response analysis of cell death in 2D plate and 3D SilkInk cultures after 5-FU exposure. The highlighted concentration indicates the dose selected for subsequent experiments. (B) Quantification of proplatelet formation across increasing 5-FU concentrations in 2D plate and 3D SilkInk cultures. (C) Platelet output across increasing 5-FU concentrations in 2D plate and 3D SilkInk cultures, expressed as fold change relative to untreated controls. Data are shown as mean ± SD. $*p < 0.05$, $**p < 0.01$, $***p < 0.001$.

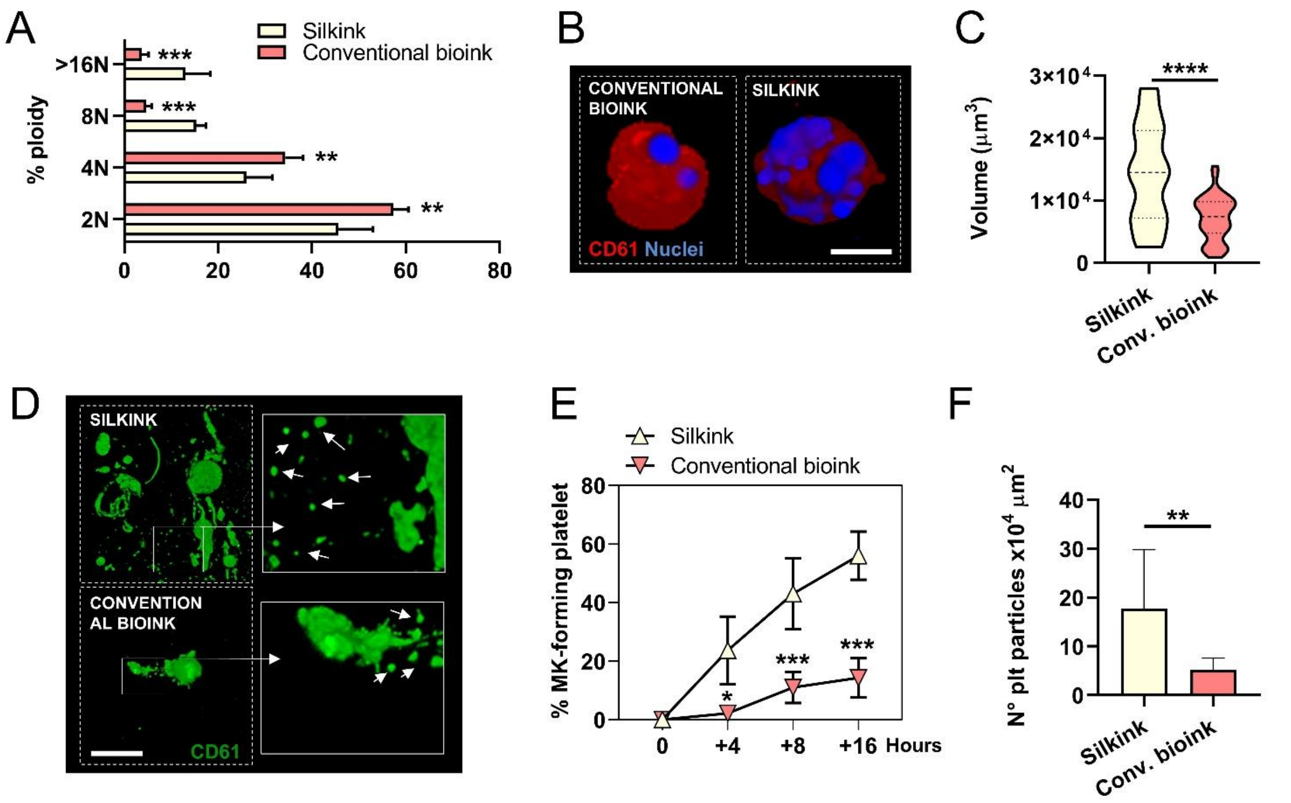


**SUPPLEMENTAL FIGURE 6. Comparison of thrombopoiesis in the 3D SilkInk with a conventional bioink.** (A) Ploidy distribution of megakaryocytes differentiated in SilkInk or in a conventional bioink. (B) Representative 3D reconstructions of CD61-positive megakaryocytes cultured in SilkInk or in a conventional bioink. Nuclei are shown in blue. Scale bar, 10 µm. (C) Quantification of megakaryocyte volume in SilkInk and conventional bioink cultures. (D) Representative confocal images showing CD61-positive proplatelets and platelet-like particles in SilkInk and conventional bioink cultures, Scale bar, 50 µm. (E) Quantification of the percentage of megakaryocytes forming proplatelets over time in SilkInk and conventional bioink cultures. (F) Quantification of the number of platelet-like particles/area ($10^4$ $\mu m^2$) in SilkInk and conventional bioink cultures. Data are shown as mean ± SD, or violin plots with median and quartiles. *$p < 0.05$, **$p < 0.01$, ***$p < 0.001$, ****$p < 0.0001$.
